# CelltrackR: an R package for fast and flexible analysis of immune cell migration data

**DOI:** 10.1101/670505

**Authors:** Inge M. N. Wortel, Katharina Dannenberg, Jeffrey C. Berry, Mark J. Miller, Johannes Textor

**Author notes:** Authors contributed equally (Mark J. Miller, Johannes Textor).

## Abstract

**Summary:** Visualization of cell migration via time-lapse microscopy has greatly advanced our understanding of the immune system. However, subtle differences in migration dynamics are easily obscured by biases and imaging artifacts. While several analysis methods have been suggested to address these issues, an integrated tool implementing them is currently lacking. Here, we present CelltrackR, an R package containing a diverse set of state-of-the-art analysis methods for (immune) cell tracks. CelltrackR supports the complete pipeline for track analysis by providing methods for data management, quality control, extracting and visualizing migration statistics, clustering tracks, and simulating cell migration.

**Availability and Implementation:** CelltrackR is an open-source package released under the GPL-2 license, and is freely available on GitHub at https://github.com/ingewortel/celltrackR.

## 1 Introduction

The ability to visualize immune cell migration using time-lapse microscopy has allowed researchers to start unraveling the cellular mechanisms underlying immunity, infection, cancer, and chronic inflammation (Moreau *et al.*, 2018), but the new data have also raised many questions. To truly understand how immune cells adjust their migration mode in different contexts, reliable quantification methods are needed.

A major challenge in extracting robust conclusions from immune cell migration data is that differences are often hard to detect, and can be obscured by imaging artifacts and biases in the analysis (Beltman *et al.*, 2009). Yet even very subtle differences in migration statistics can have large functional consequences on time scales beyond that of the imaging experiment (Textor *et al.*, 2011; Ariotti *et al.*, 2015). Although novel analysis methods and modeling approaches have been developed to deal with these issues, these are often implemented in custom-made scripts – hampering their widespread use by the community. A single, easily accessible tool integrating these different methods is currently lacking.

We here present celltrackR, an R package for the robust quantification and interpretation of immune cell migration data. Building on the powerful statistical and visualization methods already available within the R programming language (R Core Team, 2019), celltrackR supports the full workflow from visualising and quantifying cell tracks to modelling and inferring robust conclusions (Figure 1).

**Figure 1:**
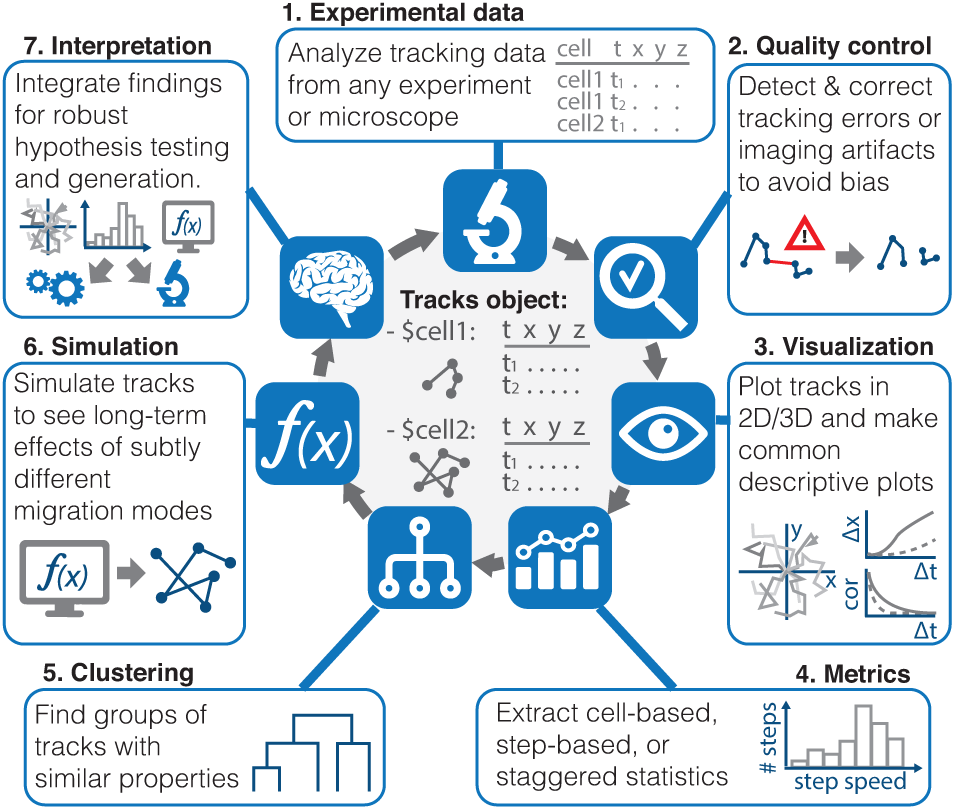
CelltrackR supports the full pipeline from cell migration data to its interpretation. The package implements a new data structure for rapid quantification of migration metrics on track datasets – the *tracks object* – as well as methods for track quality control, quantification, visualization, clustering, and simulation.

## 2 CelltrackR analysis workflow

Below, we briefly explain how celltrackR supports the cell track data analysis workflow. Detailed examples can be found in the package vignettes available via `browseVignettes(“celltrackR”)`.

### 2.1 Loading data from different experimental set-ups

Cell migration data is typically stored in the format of cell *tracks*, tables linking the position of each cell to the corresponding timepoint in time-lapse images. By working with this standardized format, CelltrackR is compatible with migration data from any experiment or microscope. It implements a special data structure for cell tracks, the *tracks object* (Figure 1). This object is a list with a coordinate matrix for each track in the dataset, and can easily be generated from a text file using the function `read.tracks.csv`. CelltrackR also supports conversion between track objects and other data structures for compatibility with custom analyses.

### 2.2 Quality control and preprocessing

CelltrackR offers several methods for track preprocessing and quality control. The `timeStep` function can be used to detect unequal time differences and missing data between subsequent time-lapse images. The `repairGaps` function fixes this issue either by splitting the problematic track, or via interpolation to more equally distributed timepoints. In general, the function `interpolateTrack` can interpolate cell positions at any timepoint of interest, which allows comparison of datasets imaged at different time resolutions. CelltrackR also implements several angle analysis tools designed to detect and deal with imaging artifacts (Beltman *et al.*, 2009). These methods are documented under ? `AngleAnalysis`.

### 2.3 Extracting and visualizing cell- or step-based statistics

Quantification of cell migration data relies on the computation of statistics such as speed, displacement, and turning angles. CelltrackR contains a range of motility statistics designed to characterise cell speed, straightness, and directionality (Mokhtari *et al.*, 2013; Gneiting and Schlather, 2004). See the documentation under ?`TrackMeasures` for details.

While it is possible to assess these statistics on tracks from individual cells, it has been shown that this “cell-based” method can introduce biases in the analysis (Beltman *et al.*, 2009). Alternatives are “step-based” (Beltman *et al.*, 2009) and “staggered” (Mokhtari *et al.*, 2013) approaches, which compute these metrics on local parts of tracks instead. CelltrackR was designed for compatibility with each of these analysis methods, allowing rapid computation of both existing and custom migration statistics in a cell-based, step-based, or staggered manner.

After track quantification, usemean square displacement plotsr, s can compare and visualize migration statistics using R’s standard statistical and visualization tools. Popular visualizations such as rose plots, mean square displacement plots, and autocorrelation plots can all be generated in this fashion and can be compared between different experiments. In addition, celltrackR implements `hotellingsTest` for an unbiased visualization and statistical analysis of subtle directionality in a dataset (Textor *et al.*, 2011).

### 2.4 Grouping and clustering tracks

The abovementioned method of extracting, visualizing, and comparing migration statistics between different datasets allows the user to perform a supervised analysis of data from different cells or experimental conditions. In addition, celltrackR also supports unsupervised track analysis. The function `clusterTracks` clusters tracks in a dataset based on one or more migration statistics using one of several popular clustering methods. In addition, the function `selectTracks` allows the user to select subsets of similar tracks by thresholding on migration statistics. These subsets can then be compared in the regular fashion.

### 2.5 Simulating immune cell migration

Since migration experiments typically measure immune cell movement over short timeframes, it can be difficult to estimate whether a subtle difference in migration statistics has any functional significance. Simulation can be a powerful tool to extrapolate experimentally observed changes in migration to their functional effects in the long run (Beauchemin *et al.*, 2007; Textor *et al.*, 2011; Ariotti *et al.*, 2015), and celltrackR supports several methods for simulating tracks. The function `brownianTrack` simulates a simple random walk, and the function `beaucheminTrack` implements a variation of this model that was designed specifically for T cells and can also simulate directionally biased motion (Beauchemin *et al.*, 2007; Textor *et al.*, 2013). In addition, `bootstrapTrack` allows the user to simulate tracks directly from migration statistics observed in a real dataset. These functions allow the user to explore long-term effects of small differences in migration statistics *in silico*, thus assisting the interpretation of immune cell migration data observed *in vitro* or *in vivo*.

## 3 Conclusion

CelltrackR implements a special data structure for migration data that allows rapid computation of a diverse array of statistics on immune cell tracks, is compatible with both cell- and step-based approaches suggested in the literature, and can easily be extended with future methods if required. The package supports several important quality controls suggested in literature and allows the user to combine track analysis, visualization, clustering, and simulation in a single platform.

## Acknowledgements

The authors thank Michael Richardson, Bernd Zin-selmeyer, Samantha Hamilton, and Gerhard Burger for their helpful feedback on the package during its beta testing.

## References

Ariotti, S., et al. (2015). Subtle CXCR3-Dependent Chemotaxis of CTLs within Infected Tissue Allows Efficient Target Localization. The Journal of Immunology, 195(11), 5285–5295.

Beauchemin, C., Dixit, N. M., and Perelson, A. S. (2007). Characterizing T Cell Movement within Lymph Nodes in the Absence of Antigen. The Journal of Immunology, 178(9), 5505–5512.

Beltman, J. B., Mareé, A. F. M., and de Boer, R. J. (2009). Analysing immune cell migration. Nature Reviews Immunology, 9(11), 789–798.

Gneiting, T. and Schlather, M. (2004). Stochastic Models That Separate Fractal Dimension and the Hurst Effect. SIAM Review, 46(2), 269–282.

Mokhtari, Z., et al. (2013). Automated Characterization and Parameter-Free Classification of Cell Tracks Based on Local Migration Behavior. PLoS ONE, 8(12), e80808.

Moreau, H. D. et al. (2018). Integrating Physical and Molecular Insights on Immune Cell Migration. Trends in Immunology, 39(8), 632–643.

R Core Team (2019). R: A Language and Environment for Statistical Computing. R Foundation for Statistical Computing, Vienna, Austria.

Textor, J., et al. (2011). Defining the quantitative limits of intravital two-photon lymphocyte tracking. Proceedings of the National Academy of Sciences, 108(30), 12401–12406.

Textor, J. et al. (2013). Analytical results on the Beauchemin model of lymphocyte migration. BMC Bioinformatics, 14(S6).

